# Oligogenic Adaptation, Soft Sweeps, and Parallel Melanic Evolution in *Drosophila melanogaster*

**DOI:** 10.1101/058008

**Authors:** Héloïse Bastide, Jeremy D. Lange, Justin B. Lack, Yassin Amir, John E. Pool

## Abstract

Unraveling the genetic architecture of adaptive phenotypic divergence is a fundamental quest in evolutionary biology. In *Drosophila melanogaster,* high-altitude melanism has evolved in separate mountain ranges in sub-Saharan Africa, potentially as an adaptation to UV intensity. We investigated the genetic basis of this melanism in three populations using a new bulk segregant analysis mapping method. Although hundreds of genes are known to affect cuticular pigmentation in *D. melanogaster,* we identified only 19 distinct QTLs from 9 mapping crosses, with several QTL peaks being shared among two or all populations. Surprisingly, we did not find wide signals of genetic differentiation (*F*_*st*_) between lightly and darkly pigmented populations at these QTLs, in spite of the pronounced phenotypic difference in pigmentation. Instead, we found small numbers of highly differentiated SNPs at the probable causative genes. A simulation analysis showed that these patterns of polymorphism are consistent with selection on standing genetic variation (leading to “soft sweeps“). Our results thus support a role for oligogenic selection on standing genetic variation in driving parallel ecological adaptation.

## Introduction

Important controversies persist regarding the process of adaptive trait evolution at the genetic level. First, phenotypic evolution may generally depend on “oligogenic” changes involving few loci or “polygenic” changes involving many loci (Bell 2009; Pritchard *et al.* 2010). The continuous nature of most phenotypic variation could imply a wealth of small effects for selection to act upon. In the presence of migration between differentially adapted populations, however, larger effects may predominate (Yeaman and Whitlock 2011). Second, the molecular properties of beneficial mutations are debated, especially the relative importance of protein-coding versus gene-regulatory changes (Hoekstra and Coyne 2007; Carroll 2008). It seems clear that positive selection acts on both types of changes, although probably favoring a greater number of noncoding variants overall (Andolfatto 2005). Still, the types of mutations acted on by selection may depend strongly upon the trait being favored (*e.g.* morphological versus physiological traits; Liao *et al.* 2010). Third, the contribution of standing genetic variation to adaptive change, relative to newly-occurring mutations, remains unresolved (Pritchard *et al.* 2010; Jensen 2014). Selection favoring a preexisting genetic variant may bring multiple haplotypes to high frequency, leading to a “soft selective sweep” that may yield much different patterns of genetic variation than a classic “hard sweep” (Pennings and Hermisson 2006). A final question concerns the genetic predictability of adaptive trait evolution - when the same phenotype arises in two or more populations or species, how often does natural selection act on the same genes or even the same variants? Resolving these biologically important questions will require adding further empirical case studies of the genetic basis of adaptive evolution.

Coloration has broad adaptive significance in survival and reproduction (Majerus 1998), making it an attractive target for genetic study. In most animals, melanin is synthesized by a small number of proteins whose patterning and sexual differentiation may be controlled by a myriad of transcription factors (Kronforst *et al.* 2012). In the fruit fly *Drosophila melanogaster,* wherein the melanin-synthesis pathway is relatively clearly defined (Wittkopp *et al.* 2003; Massey and Wittkopp 2016), a recent study identified 28 trans-regulators of pigmentation (Rogers *et al.* 2014), and mutational screens have identified more than four hundred genes that may impact body color (flybase.org as of April 2016). And yet, genome-wide association studies (GWAS) of female abdominal pigmentation variation in four temperate populations all revealed major effects of the two melanin-synthesis genes *tan* and *ebony* and the transcription factor *bric-a-brac 1 (bab1),* although other minor effect genes were also detected (Bastide *et al.* 2013; Endler *et al.* 2015; Dembeck *et al.* 2015).

Pigmentation displays strong geographic trends in *D. melanogaster.* Several studies have reported a trend of darker cuticle at high latitudes in non-African populations (David *et al.* 1985; Das 1995; Munjal *et al.* 1997; Telonis-Scott *et al.* 2011), and an enrichment of pigmentation genes in genome windows differentiating northern and southern populations was detected in a genome-wide selection scan in Australia (Reinhardt *et al.* 2014), paralleling the cuticular pigmentation cline found on this continent (Telonis-Scott *et al.* 2011). In tropical Africa, which harbors the ancestral range of the species (David and Capy 1988; Pool *et al.* 2012), unusually dark populations have been discovered in different mountain ranges (Pool and Aquadro 2007; Bastide *et al.* 2014). Overall, the pigmentation of African *D. melanogaster* is best predicted by UV intensity, offering a plausible selective agent to drive the recurrent evolution of melanism (Bastide *et al.* 2014). In one population (Uganda), a haplotype carrying a series of causative cis-regulatory mutations at *ebony* was a major contributor to melanism and showed evidence of a strong partial selective sweep (Pool and Aquadro 2007; Rebeiz *et al.* 2009). However, no unbiased genome-wide search for loci underlying this parallel color evolution has been conducted.

Here, we investigate the genetic basis of melanic flies in three populations from Ethiopia, Cameroon and Uganda, presenting the first application of a bulk segregant analysis (BSA) approach designed for *Drosophila* (Pool 2016). We then compared the level of genetic differentiation at the pigmentation-associated regions between each of the melanic populations and a lightly pigmented population from Zambia. Many of the identified QTLs contained major melanin-synthesis genes with varying genetic differentiation between the light and dark populations. Analysis of demography and differentiation at these genes supported a role of natural selection on standing genetic variation in parallel melanic adaptation.

## Materials and Methods

### Natural populations investigated

The populations in the present study were all studied by Bastide *et al.* (2014), where a number of individuals coming from 30 natural populations of various latitudes and altitudes were scored for body pigmentation. Among those, three Afrotropical populations showed an outstanding dark pigmentation and were used to found experimental crosses: Fiche, Ethiopia (EF, 9.81° N, 38.63° E, alt. 3070 m) showing the most extreme phenotype with the entire body of the fly strongly melanized; Oku, Cameroon (CO, 6.25° N, 10.43° E, alt. 2169 m); and Namulonge, Uganda (UG, 0.53° N, 32.60° E, alt. 1134 m). In addition, a population from Siavonga, Zambia (ZI, 16.54° S, 28.72° E, alt. 530 m) was chosen as the reference light population against which all of the dark populations were subsequently crossed (Figure 1).

**Figure 1.**
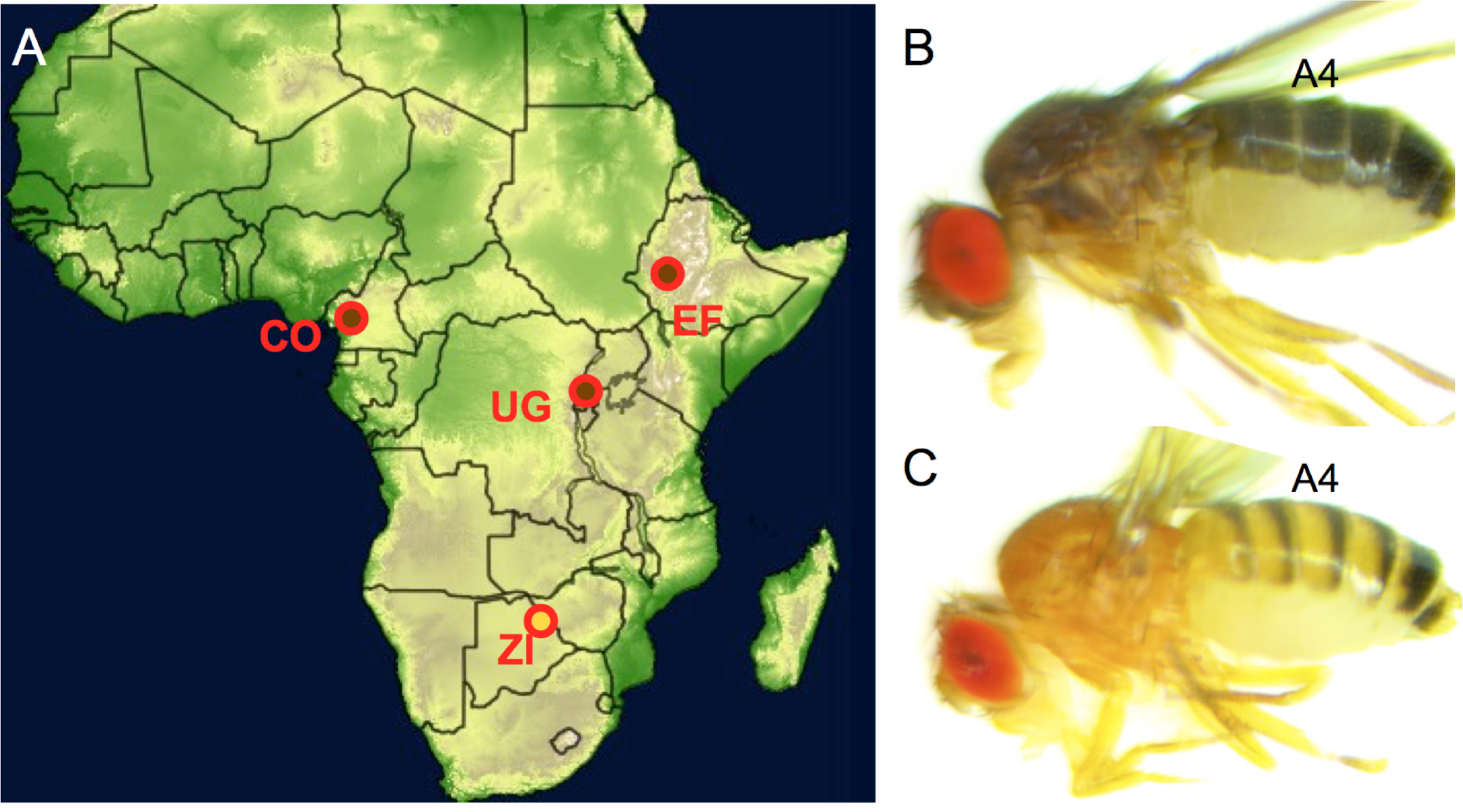
Populations sampled and studied phenotypes. (A) Several lines of each melanic population (brown circle; CO = Cameroon, EF = Ethiopia, UG = Uganda) were separately crossed with homokaryotypic lines from a lightly pigmented population (yellow circle; ZI = Zambia). (B and C) Pigmentation phenotypes in Ethiopia (B) and Zambia (C) showing the fourth abdominal segment that was analyzed for mapping.

### Choice ofparental lines for outbreeding crosses

We set 4, 3 and 2 experimental crosses each seeded with F1 flies produced from a cross between a darkly pigmented EF, CO or UG line and a lightly pigmented ZI line, respectively (Table S1). Each melanic line was only used in one mapping cross, and all had previously been inbred for 8 generations. To choose the parental lines, we tested lines from each population for the presence of eight common chromosomal inversions by PCR reaction using primers and amplification conditions from Lack *et al.* (2016) to avoid seeding a cross with heterokaryotypic flies unable to recombine near the inversion. These included six autosomal cosmopolitan inversions *(In(2L)t, In(2R)NS, In(3L)P, In(3R)P, In(3R)K,* and *In(3R)Mo)* and two sub-Saharan inversions on the X chromosome *(In(1)A* and *In(1)Be.* We also used the pigmentation score data of the female fourth abdominal segment that we generated in Bastide *et al.* (2014) to select darker lines for each melanic population (in conjunction with other phenotypes of interest).

### Experimental design for the generation of outcrossed flies

We followed the experimental design described in Pool (2016). For each cross, we conducted reciprocal crosses between 8 melanic strain flies and 8 ZI strain flies, independently. From each of these two crosses, 125 random F1 males and 125 random F1 females were mixed altogether (N = 500) and allowed to evolve for several non-overlapping generations at a constant population size of ~1,500 individuals in 28 × 14 × 15 cm plastic cages, each provided with 14 vials with standard *Drosophila* medium (containing molasses, corn meal, yeast, agar, and antimicrobial agents) and kept at ~20° C. Each generation, adults were allowed to lay eggs on the food for one week and then discarded, and food vials were replaced every 3 weeks when adult flies in the cage are 7-10 day old. After 20 generations (allowing a large number of unique recombination events to take place), we allowed adult flies to lay eggs on fresh food for 2 days before replacing the vials and waiting for F20 flies to emerge. We then visually phenotyped 3-5 day old females (*N* = 600) for pigmentation on the fourth abdominal segment under CO2 anesthesia, focusing on either pigmentation intensity near the anterior margin (A4 background; crosses EB1, EB2 and UB1) or the width of the posterior black stripe (A4 stripe width; crosses CS1, CS2, CS3, ES1, ES2 and US1) as in Bastide et al. (2014). For each cross, flies were sorted into the 10% darkest (*N* = 60) and the 10% lightest (N = 60) females.

### Preparation of genome libraries

Genome libraries were independently prepared for the parental lines and the four F_20_ pigmentation subgroups for each cross. For each library, genomic DNA was extracted from a pool of 30 females by chloroform extraction and ethanol precipitation. DNA was then fragmented using the Bioruptor sonicator (Diagenode), and paired-end libraries with approximately 300 bp inserts were prepared using NEBNext DNA Library Prep Reagent Set for Illumina (New England Biolabs no. E6000L). Library concentration and quality were assessed using an Agilent 2100 Bioanalyzer (Agilent Technologies, Inc.) and were sequenced at the UW-Madison Biotechnology Center on the Illumina HiSeq 2000 platform with 100 bp paired read lengths.

### Alignment of the raw sequences

Reads were mapped to the *D. melanogaster* reference genome (release 5.57), using default parameters in BWA v0.6.2-r126 (Li and Durbin 2009). The BAM files were remapped with Stampy v1.0.21 (Lunter and Goodson 2011) and the reads were filtered for a mapping quality of 20 and for proper pairs with samtools v0.1.18 (Li *et al.* 2009). BAM files were cleaned by removing unmapped reads and sorted by coordinate, and PCR duplicates were marked using Picard v1.109 (http://picard.sourceforge.net). Alignment around indels was then improved using GATK v3.2 (McKenna *et al.* 2010; DePristo *et al.* 2011). Sequencing depth obtained for each mapping population sequencing is given in Table S1.

### Genome mapping of pigmentation genes using ancestry analysis

For each cross, we used the PoPoolation2 ver. 1.201 software package (Kofler *et al.* 2011) to generate a synchronized mpileup file for the two parental genomes and the pigmentation-sorted pools aligned to the *D. melanogaster* reference. For each biallelic SNP, an ancestry difference value (*ad*) summarized the difference in parental strain ancestry between the high and low phenotypic pools. With respect to the melanic parental line allele, *a_d_* was estimated as

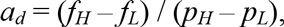

where *p_H_* is the frequency of the major allele in the melanic parental line,*p_L_* the frequency of the melanic allele in the non-melanic parental line, *fH* the frequency of the melanic allele in the F20 dark subgroup, and *f_L_* is the frequency of the melanic allele in the F_20_ light subgroup. Only sites with parental strain frequency difference*p_H_—p_L_* ≥ 0.25 were considered. The five chromosomal arms (X, 2L, 2R, 3L and 3R) were subdivided into 2728, 3131, 2357, 2956 and 2935 windows, each of roughly 8.4 kb on average, whose boundaries were determined according SNP density in ZI genomes (Lack *et al.* 2015). For each cross, ancestry difference values were averaged across qualifying SNPs for each window. To summarize trends across windows, we performed a simple smoothing of empirical and simulated ancestry difference values. We weighted the focal window's value with a factor of 5, and gave descending weights of 4, 3, 2, and 1 for the 4 windows on each side.

QTL mapping was performed using Simulation-based Inference for Bulk Segregation Analysis Mapping (SIBSAM; Pool 2016). Unlike BSA in yeast, where millions of segregants can be generated, BSA using *Drosophila* may often generate overlapping QTL peaks, which most BSA mapping approaches are not designed to account for. SIBSAM therefore analyzes both primary QTL peaks (the maximum value in an interval of continuously positive *a_d_*) and secondary QTL peaks that may flank them (Pool 2016). Simulations are conducted based on the full experimental process (recombination in multiple males and females for multiple generations), selection on phenotype in the final generation (which is based on diploid genotype at each QTL and environmental/measurement variance), and the sampling of sequence reads to obtain *a_d_.* SIBSAM involves a three phase inference process that results in estimates of the significance, location, and strength of QTLs. First, simulations under the null model (no true QTLs) are conducted in order to assess the significance of primary peaks. Second, simulations with a single QTL are conducted, which result in estimates of QTL effect size (proportion of phenotypic variance explained), confidence intervals for that quantity and for genomic location of the true QTL, and the significance of secondary QTL peaks. If significant secondary peaks are present, simulations of multiple linked QTLs are conducted in order to refine the strength and location estimates of the primary and secondary peaks (Pool 2016).

### Scan of genetic differentiation

In order to test whether significant QTLs contained highly differentiated windows between the two populations from which the parental lines were drawn, we estimated *F_ST_* for these windows between Zambia and the melanic population in question. Genomes from ZI (197), CO (10), EF (68), and Uganda (40) were obtained from the *Drosophila* Genome Nexus (Lack *et al.* 2015). The Uganda population consisted of a pool of lines from Rwanda and Uganda that show minimal genetic differentiation *(F_ST_* = 0.015; Pool *et al.* 2012) and similarly dark pigmentation (Bastide *et al.* 2014). For each window, we also estimated the quantile (Q) of the window *F_ST_* relative to the empirical distribution of *F_ST_* of all windows per chromosomal arm, where *Q* denotes the proportion of windows with an equal or greater *FST* value. Strong selective sweeps from new mutations are expected to produce high window-based *FST* values due to hitchhiking effect, but such outliers may not include soft sweeps (selection on existing alleles that already occur on multiple haplotypes). Consequently, we also estimated the maximum SNP-based *F_ST_* in each window and the quantile of this value according to the empirical distribution of this estimate for all windows per chromosomal arm.

### Estimating a neutral null model for the Ethiopian population

The EF population harbors flies with some of the darkest phenotypes among all *D. melanogaster* populations (Bastide *et al.* 2014). Because the melanin-synthesis gene *ebony* has previously been shown to be implicated in melanism in non-Ethiopian African populations (Pool and Aquadro 2007; Rebeiz *et al.* 2009) and was also found in our mapping analyses in EF (see Results), we compared its observed polymorphism to that expected under different selection scenarios. To estimate a neutral model consistent with the genetic diversity of this population, we used the allele frequency spectrum (AFS) from 105,715 SNPs from the EF, Rwanda (RG) and ZI populations, respectively, falling within autosomal short intronic segments (< 86 bps, with 16 and 6 bps removed from the intron start and end, respectively). These sites are presumably neutral (Halligan and Keightley 2006), although they can be affected by selection at linked sites. We then fit our observed AFS of the three populations to the implemented “out of Africa” demographic model using a diffusion approximation approach implemented in the δαδ*l* ver. 1.7 software package (Gutenkunst *et al.* 2009). This model (Figure S1) implies an instantaneous growth of the ancestral population (ZI) prior to an initial split, with a bottleneck taking place in the ancestor of the RG and EF populations. A second split between the two derived populations takes place followed by a bottleneck and a growth phase in each descendant population, with ongoing migration occurring among all populations. Parameters were optimized using δαδι and several runs were conducted to confirm that convergent parameter estimates were obtained. We then used non-parametric bootstrapping to infer parameter uncertainties. For this, we generated 100 bootstrapped AFS from the empirical data, and parameter standard deviations were estimated using the Godambe Information Matrix (GIM) approach as implemented in δαδι version 1.7. Ancestral effective population size *(N_e_)* was inferred by dividing δαδι-estimated Watterson’s theta (*θ*) over 4 times the mutation rate, using an estimate from *D. melanogaster* of 3.27 × 10^−9^ (Schrider *et al.* 2013). Divergence times (in years) were then estimated assuming 15 generations per year (Pool 2015).

### Simulations

We used *msms* (Ewing and Hermisson 2010) to simulate a region motivated by the *ebony* locus.

The optimized demographic parameters from the SaSi analysis used in these simulations were converted from units of *2N_e_* to *4N_e_* generations. Recombination rates were based on the local estimate by Comeron *et al.* (2012). The empirical data that we sought to emulate came from a 5001 base pair window centered on the most differentiated SNP. At this site, the empirical data included 33 genomes from Ethiopia EF and 189 from Zambia ZI. Our simulations matched these sample sizes while also simulating a third population, representing Rwanda RG, which was present in the demographic modeling and simulations, but not included in the analysis of simulated data. Examining a range of scenarios with and without positive selection, we examined the propensity of each simulation to generate the disparate window *FST* and maximum SNP *FST* values observed between the EF and ZI samples at *ebony.*

We first ran 10,000 neutral simulations to test whether the values observed at *ebony* were unexpected in the absence of selection. We then explored a range of scenarios with positive selection limited to the Ethiopian population. Here, we varied the selection coefficient (*s*) and the frequency of the favored allele at the start of the sweep. In these simulations, we explored the hypothesis that the most differentiated SNP observed at *ebony* was a beneficial mutation (in Ethiopia only), and we set the target of selection to be the middle of the simulated 5001 base pair locus. We simulated 10 different selection strengths and 6 starting beneficial allele frequencies for a total of 60 scenarios of selection. The 10 selection coefficients we studied were 0.00005, 0.000075, 0.0001, 0.0025, 0.005, 0.0075, 0.001, 0.0025, 0.005, and 0.0075. The 6 starting beneficial allele frequencies were 1/2N_*e*_, 0.005, 0.01, 0.025, 0.05, and 0.1.

In the empirical data, 5 out of 189 Zambia genomes contained the hypothesized beneficial variant, while 29 of 33 Ethiopian genomes carried it. In these simulations, our primary interest was to ask which models could produce this strong SNP frequency difference while generating a window *F_ST_* as low as that observed at *ebony.* In order to emulate these SNP frequencies in simulations with selection, we employed a sub-sampling strategy. We initially simulated larger samples of chromosomes for the Zambia (n=250) and Ethiopia (n=50) populations, then pared them down to 189 and 33 chromosomes, respectively. Sub-sampling was then pursued to achieve a nontrivial probability of a simulation matching our frequency criteria. Specifically, we sub-sampled to ensure that the Ethiopia population contained exactly 29 chromosomes with the beneficial allele and 4 without, while the Zambia population contained exactly 5 chromosomes matching the allele favored in Ethiopia. Simulations lacking sufficient numbers of both alleles in either population to achieve these subsamples were rejected. Thus, scenarios that consistently fixed the beneficial allele in Ethiopia or else failed to elevate its frequency yielded low acceptance rates. Although we used the-SFC flag of *msms* to condition on the beneficial allele's persistence in the Ethiopian population at the time of sampling, simulations could still be rejected if this allele was unsampled or sampled at low frequency. For each selection scenario, we ran a maximum of 2,500,000 simulations in an attempt to get 1,000 accepted simulations.

We also filtered simulations to clearly distinguish between the predictions of hard sweeps and soft sweeps. For the case of an initial beneficial allele frequency of 1/2N_e_ (corresponding to new mutations), only hard sweeps could be generated. For all other initial frequencies, we required at least two unique copies of the beneficial allele to be present at sampling, in order to study soft sweeps specifically.

## Results

### Large effect loci underlie melanism in the three populations

We used a newly-developed BSA method (Pool 2016) to map pigmentation QTLs (Figures 2, S2 and S3). We mapped two pigmentation traits (A4 abdominal background color and A4 stripe width) in darkly pigmented populations from three African populations of *D. melanogaster* (Cameroon, Ethiopia, Uganda). A total of 35 significant QTL peaks were found in the nine crosses (Table S2). Effect sizes ranged from 6.29% to 36.39% for primary peaks and from 6.72 to 19.20% for secondary peaks. Based on overlapping genomic confidence intervals, these can be summarized into 19 major QTLs, ranging from 10 kb to 3 Mb long, of which 12 were unique to different single crosses (Table 1). Notably, even when the same trait was investigated in the same population, QTLs often differed considerably between independent crosses. Many of these QTL peaks are tall enough that we expect very high power to detect them (Table S2; Pool 2016), suggesting that discordant results are unlikely to result solely from a randomly detected subset of shared QTLs (see Discussion).

**Table 1.**
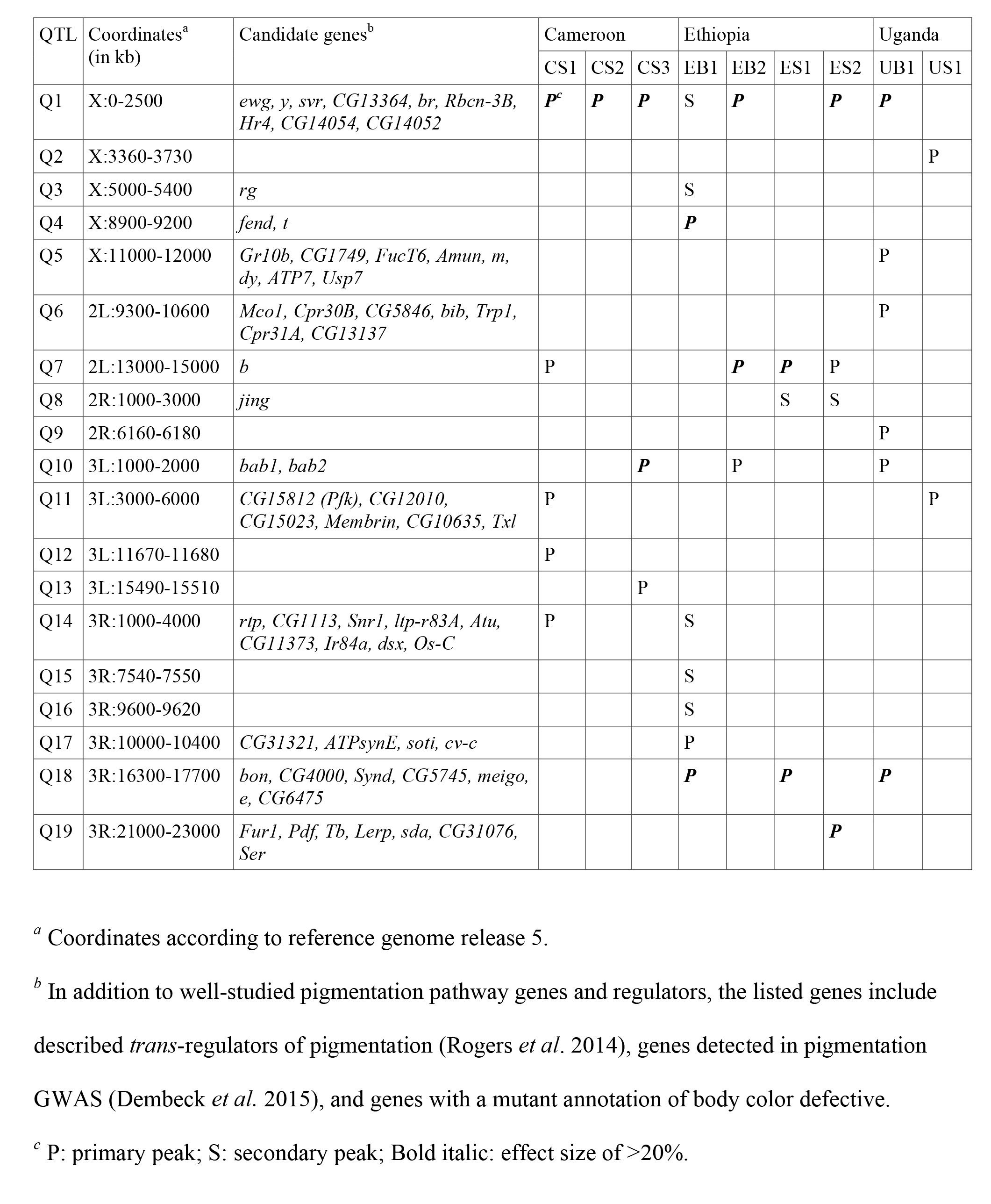
Quantitative trait loci (QTLs) underlying melanic evolution identified from nine crosses between three high-altitude and one low-altitude Sub-Saharan populations of *D. melanogaster.*

| QTL | Coordinates<sup>a</sup><br>(in kb) | Candidate genes<sup>b</sup> | Cameroon |  |  | Ethiopia |  |  |  | Uganda |  |
| --- | --- | --- | --- | --- | --- | --- | --- | --- | --- | --- | --- |
|  |  |  | CS1 | CS2 | CS3 | EB1 | EB2 | ES1 | ES2 | UB1 | US1 |
| Q1 | X:0-2500 | *ewg, y, svr, CG13364, br, Rbcn-3B, Hr4, CG14054, CG14052* | **P<sup>c</sup>** | **P** | **P** | S | **P** |  | **P** | **P** |  |
| Q2 | X:3360-3730 |  |  |  |  |  |  |  |  |  | P |
| Q3 | X:5000-5400 | *rg* |  |  |  | S |  |  |  |  |  |
| Q4 | X:8900-9200 | *fend, t* |  |  |  | **P** |  |  |  |  |  |
| Q5 | X:11000-12000 | *Gr10b, CG1749, FucT6, Amun, m, dy, ATP7, Usp7* |  |  |  |  |  |  |  | P |  |
| Q6 | 2L:9300-10600 | *Mco1, Cpr30B, CG5846, bib, Trp1, Cpr31A, CG13137* |  |  |  |  |  |  |  | P |  |
| Q7 | 2L:13000-15000 | *b* | P |  |  |  | **P** | **P** | P |  |  |
| Q8 | 2R:1000-3000 | *jing* |  |  |  |  |  | S | S |  |  |
| Q9 | 2R:6160-6180 |  |  |  |  |  |  |  |  | P |  |
| Q10 | 3L:1000-2000 | *bab1, bab2* |  |  | **P** |  | P |  |  | P |  |
| Q11 | 3L:3000-6000 | *CG15812 (Pfk), CG12010, CG15023, Membrin, CG10635, Tx1* | P |  |  |  |  |  |  |  | P |
| Q12 | 3L:11670-11680 |  | P |  |  |  |  |  |  |  |  |
| Q13 | 3L:15490-15510 |  |  |  | P |  |  |  |  |  |  |
| Q14 | 3R:1000-4000 | *rtp, CG1113, Snr1, ltp-r83A, Atu, CG11373, Ir84a, dsx, Os-C* | P |  |  | S |  |  |  |  |  |
| Q15 | 3R:7540-7550 |  |  |  |  | S |  |  |  |  |  |
| Q16 | 3R:9600-9620 |  |  |  |  | S |  |  |  |  |  |
| Q17 | 3R:10000-10400 | *CG31321, ATPsynE, soti, cv-c* |  |  |  | P |  |  |  |  |  |
| Q18 | 3R:16300-17700 | *bon, CG4000, Synd, CG5745, meigo, e, CG6475* |  |  |  | **P** |  | **P** |  | **P** |  |
| Q19 | 3R:21000-23000 | *Fur1, Pdf, Tb, Lerp, sda, CG31076, Ser* |  |  |  |  |  |  | **P** |  |  |

<sup>a</sup> Coordinates according to reference genome release 5.
<sup>b</sup> In addition to well-studied pigmentation pathway genes and regulators, the listed genes include described *trans*-regulators of pigmentation (Rogers *et al.* 2014), genes detected in pigmentation GWAS (Dembeck *et al.* 2015), and genes with a mutant annotation of body color defective.
<sup>c</sup> P: primary peak; S: secondary peak; Bold italic: effect size of >20%.

**Figure 2.**
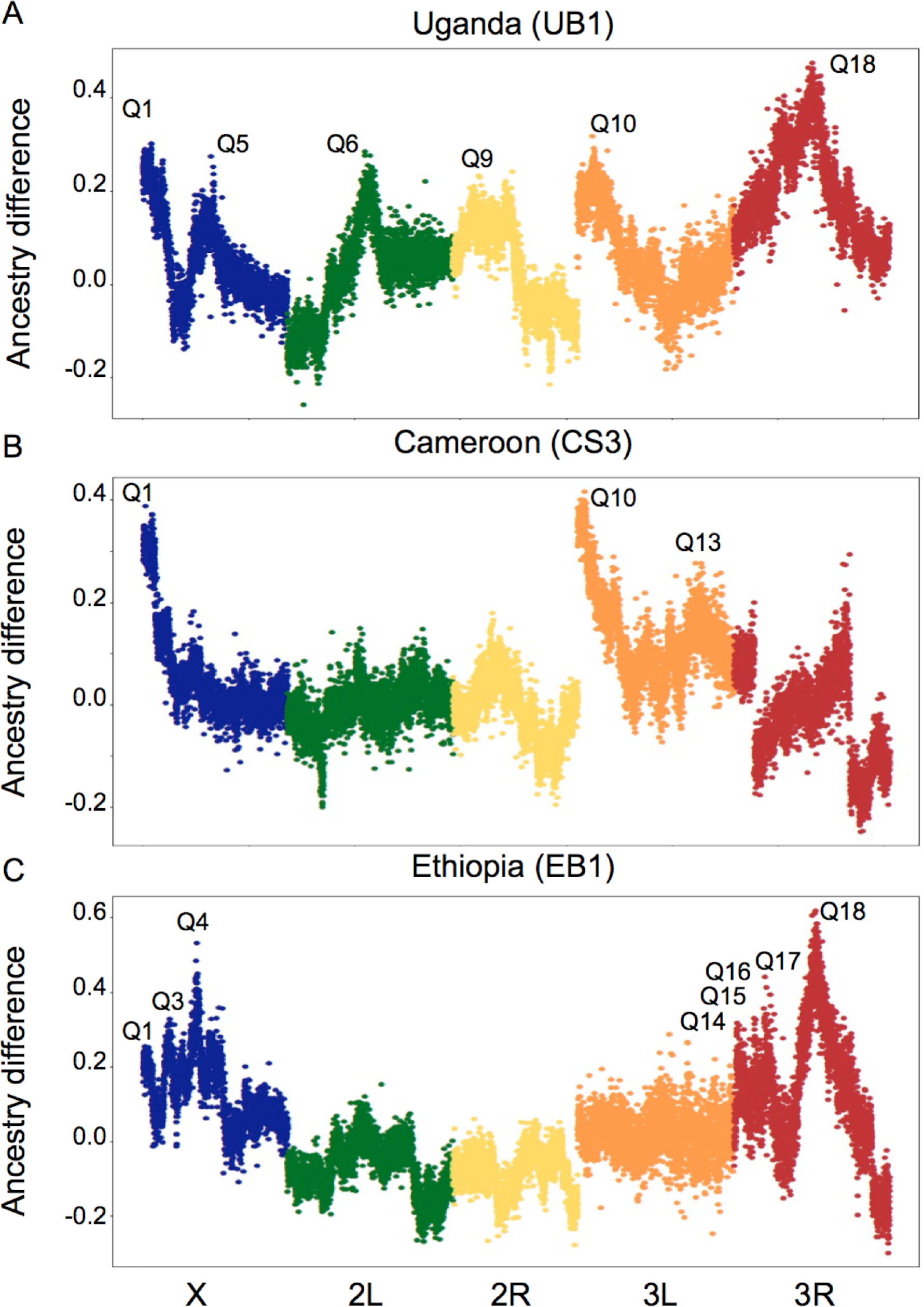
Ancestry difference plots showing relative proportions of the melanic parental population allele across five colored chromosomal arms (X, 2L, 2R, 3L, 3R) in crosses involving three melanic populations: (A) Uganda (UB1), (B) Cameroon (CS3) and (C) Ethiopia (EB1). This raw mapping surface is an input for SIBSAM. QTL names are according to Table 1. Discontinuities in the Cameroon plot’s chromosome arm 3R reflect the presence of *In(3R)K* in both parental strains, an inversion that is nearly fixed in the CO sample.

Among the common QTL peak regions, one (Q1) was shared among seven crosses while another was shared among four crosses. The former region consists of the 2.5 Mb telomeric end of chromosome X containing the important melanin-synthesis gene *yellow.* The second most common QTL region (Q7) extends for 2 Mb and contains another melanin-synthesis gene, *black,* and it was shared among three Ethiopian and one Cameroonian crosses. These two genes have not been detected in pigmentation analyses within cosmopolitan populations. While *yellow* has been implicated in the evolution of pigmentation between multiple *Drosophila* species (Wittkopp *et al.* 2002; Jeong *et al.* 2006; Ordway *et al.* 2014), to our knowledge *black* has never been found to underlie *Drosophila* pigmentation evolution (Massey and Wittkopp 2016). The enzyme coded by *black* converts uracil to ×-alanine. The enzyme coded by *ebony* conjugates this to dopamine to form N-ß-alanyl-dopamine (NBAD), a precursor of light pigmentation. The enzyme coded by *tan* converts NBAD back to dopamine, a precursor of dark pigmentation (Wittkopp *et al.* 2003). *black* thus constitutes an integral part of the *tan-ebony* loop, the most evolutionary labile part of the melanin synthesis network in *Drosophila*.

Two other QTL peak regions were each shared among three crosses. One (Q10) included the transcription factor *bab1* and was shared by one cross from each of the three melanic populations. The other region (Q18) was shared by two crosses from Ethiopia and one cross from Uganda, and includes the melanin-synthesis gene *ebony.* In Uganda, a strong effect of *ebony* on pigmentation has previously been illustrated (Pool and Aquadro 2007; Rebeiz *et al.* 2009), providing a “positive control” for our mapping method. Both *bab1* and *ebony* are associated with pigmentation variation in cosmopolitan populations (Kopp *et al.* 2003; Bickel *et al.* 2011; Bastide *et al.* 2013; Rogers *et al.* 2013; Endler *et al.* 2015; Dembeck *et al.* 2015; Johnson *et al.* 2015; Miyagi *et al.* 2015). The remaining peaks did not include other major melanin-synthesis enzymes (Table 1) with an exception in a single Ethiopian cross where a strong primary peak (Q4) included *tan.*

Taken together, in spite of the genetic complexity of abdominal pigmentation some common bases may persist. For example, major pigmentation loci *(yellow*, *black, bab1, ebony* and *tan)* are within Ethiopia QTLs, with some shared with both Uganda and Cameroon *(yellow* and *bab1),* some with only Cameroon *(black)* or only Uganda *(ebony),* and one that is unique to one cross of this population (tan). Our genome mapping also included other 59 genes that are known, mostly from mutational screens, to cause changes in body coloration (Table 1). None of these have been suggested before to affect pigmentation variation within or between species (Massey and Wittkopp 2016).

### Low genetic differentiation at pigmentation-associated loci in Ethiopia

In order to investigate whether pigmentation-associated peaks harbor genes that have been strongly differentiated between the lowland ZI population and each of the melanic populations, we estimated their differences in allele frequencies using *FST* for windows that have been used in the ancestry analyses (Tables S3 and S4). Consistent with previous findings of an incomplete selective sweep at the gene *ebony* in Uganda (Pool and Aquadro 2007; Rebeiz *et al.* 2009), window *F_ST_* between Uganda and Zambia detected a moderate peak around this locus, although these windows did not fall within the most extreme 5% *(i.e. Q* > 0.05; Figure 3A). Among the remaining six QTLs in this population, three contained windows with elevated differentiation, including for the X-linked melanin-synthesis gene *yellow* (Q1; *Q* < 0.001; Figure S4A), although genetic differentiation tends to be broadly elevated across this low recombination telomeric region. Elevated *F_ST_* was also detected for a window upstream of the two transcription factor paralogs *bric-a-brac* (Q10; *Q* < 0.05; Figure 3A), and Q11 containing two highly differentiated genes that may affect body color: *CG15812 (piefke)* and *CG15023* (Mummery-Widmer *et al.* 2009; *Q* < 0.01; Figure S5A).

**Figure 3.**
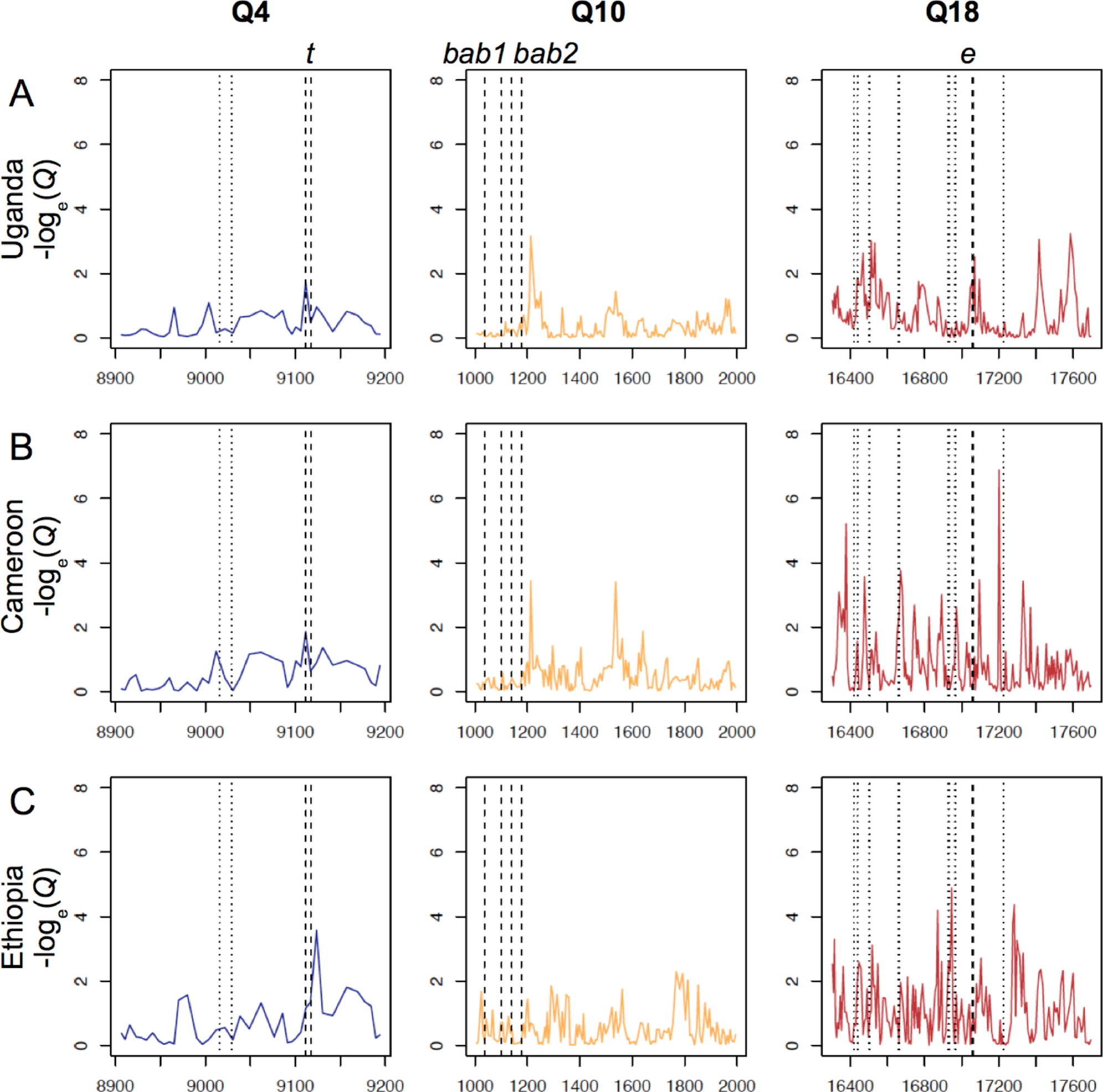
Window-based genetic differentiation *(FST)* in quantiles (Q) between a lightly pigmented population (Zambia) and three melanic populations: (A) Uganda, (B) Cameroon and (C) Ethiopia at three pigmentation-associated QTLs. Dotted lines refer to boundaries of pigmentation candidate genes: *tan* (t), *bric-a-brac (babl* and *bab2*), and *ebony* (e). Coordinates are given in kb with respect to release 5 of the *D. melanogaster* genome. In many cases, strong window genetic differentiation was not observed at pigmentation genes within large-effect QTLs.

In Cameroon, significant differentiation was found within five QTLs detected in this population, including windows similar to that found in Uganda within Q1 (*Q* < 0.001; Figure S4B), Q10 (*Q* < 0.05; Figure 3B) and Q11 (*Q* < 0.05; Figure S5B). Three peaks of differentiation (Q < 0.05) were found at Q7 surrounding the melanin-synthesis gene *black* but no peak at the gene itself (Figure S4B). None of these peaks contain any known pigmentation-related genes.

In Ethiopia, strong differentiation was found in all but 2 (Q10 and Q16) of its 12 QTLs, but in general these peaks did not fall near pigmentation candidate genes. The four melanin-synthesis genes (Figures 3C and S4C) in the QTLs of this population did not show strong signals of window genetic differentiation, including *ebony* in spite of the major effect of this QTL in the cross with the darkest phenotype. A single exception was the gene *rugose* which may affect body color (Bateman 1950) at Q3 (Q < 0.05; Figure S5C), but selection at this gene might also relate to the evolution of other phenotypes in this population.

### Strongly differentiated SNPs at candidate pigmentation genes in Ethiopia

Window-based *F_ST_* estimates are strongly influenced by the specific model of positive selection, with soft and/or incomplete sweeps potentially producing different degrees of window genetic differentiation than classic hard/complete sweeps (Lange and Pool 2016). We therefore also investigated *F_ST_* values for individual SNPs at four candidate genes within QTLs *(black, ebony, tan,* and *yellow)* for Ethiopia versus Zambia. In spite of the lack of strong window *F_ST_* signal cited above, all four of these genes had one or more individual SNPs with strong differentiation, yet without any strong pattern of differentiation at linked SNPs (Figure 4). Since individual SNP *F_ST_* may be more vulnerable to random variance when sample sizes are small, we note that for the SNPs described below, or sample sizes range from 33 to 55 genomes for Ethiopia, and from 188 to 191 for Zambia.

**Figure 4.**
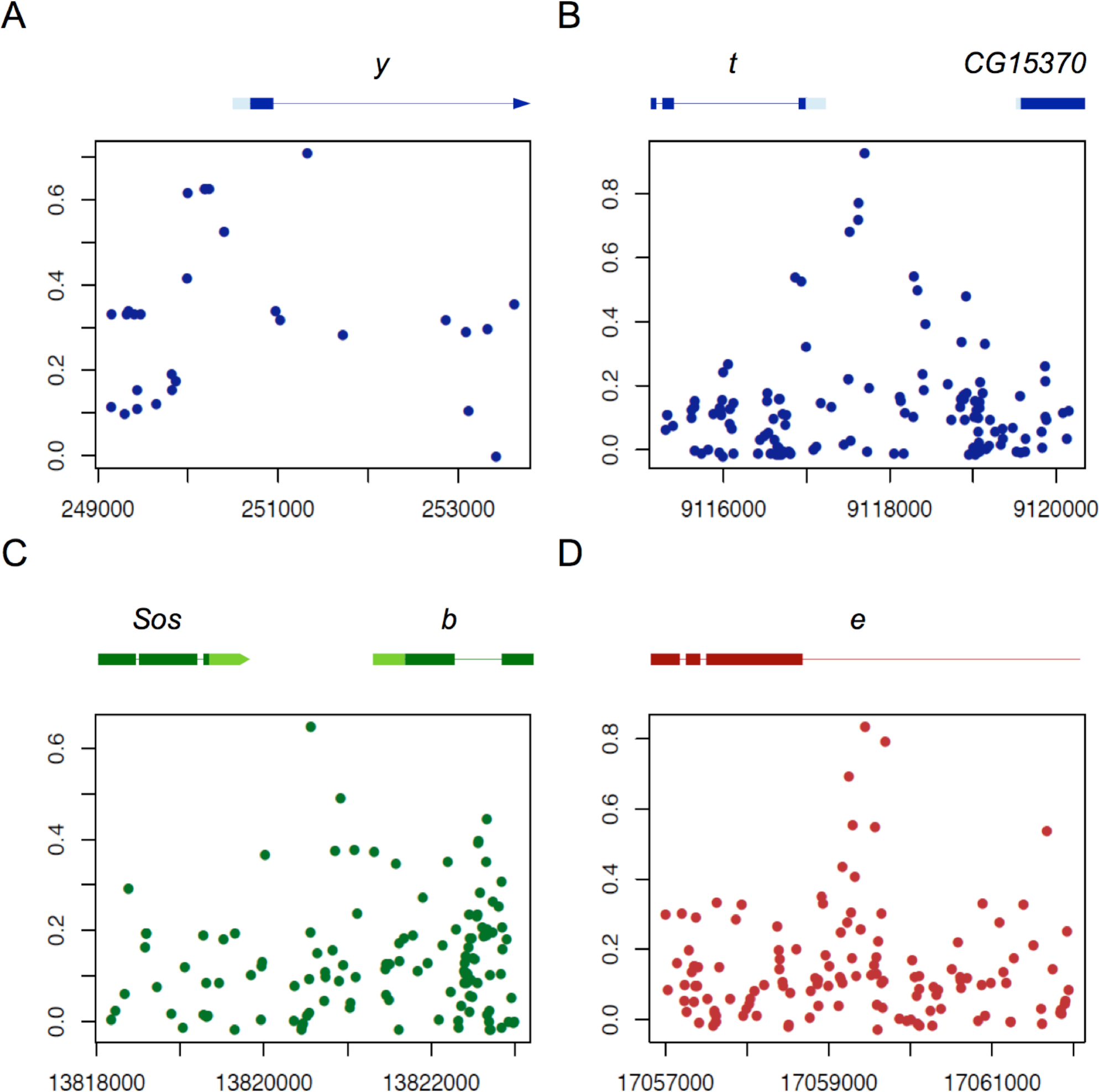
SNP-based genetic differentiation *(F_ST_*) estimates between lightly pigmented Zambia and darkly pigmented Ethiopia populations at four melanin-synthesis enzyme genes: (A) *yellow* (*y*), (B) *tan* (*t*), (C) *black* (*b*) and (D) *ebony* (*e*). Each plot represents 5-kb window centered on the most differentiated SNP for each gene. Lightly colored boxes refer to genes 5′ and 3′ UTRs, darkly colored boxes refer to exons, and lines refer to introns.

At *yellow* (Figure 4A), the most differentiated SNP (X:251,323; coordinates with respect to release 5 of the *D. melanogaster* reference genome) fell in the first intron. Although this region is not known to affect abdominal pigmentation in cosmopolitan *D. melanogaster,* in *D. pseudoobscura* and *D. virilis,* two species that are completely dark and whose phenotype resembles the dark Ethiopian phenotype, intronic enhancers affect body pigmentation (Kalay and Wittkopp 2010).

At *tan* (Figure 4B), the most differentiated SNP (X:9,117,695) did not fall in the male-specific-enhancer (t_MSE) which had often been detected in within-species studies focusing on the last abdominal segments in *Drosophila* (Bastide *et al.* 2013; Dembeck *et al.* 2015; Yassin *et al.* 2016) rather than non-sexually differentiated abdominal segments like this one. Instead, the most differentiated SNP fell 400 bp upstream of the beginning of the gene in a binding site of the transcription factor *dorsal* (Drosophila modENCODE Consortium 2010), which is involved in the melanization defense response (Bettencourt *et al.* 2004). This SNP is tightly linked to a short indel 7 bp away. Although SNPs in the first intron of *tan* are associated with dark pigmentation in the *virilis* species group, enhancers in the proximal upstream region are yet to be investigated.

At *black* (Figure 4C), the most differentiated SNP (2L:13,820,561) falls 700 bp upstream the beginning of the gene in a region that was shown to bind with the homeotic transcription factor *prd* (Drosophila modENCODE Consortium 2010). The enhancers of *black* affecting pigmentation in different body parts are yet to be studied.

At *ebony* (Figure 4D), the most differentiated SNP (3R:17,059,445) in Ethiopia falls within the first intron, which co-regulates *ebony* expression in the abdomen together with an upstream enhancer known as the core abdominal cis-regulatory element (e_abdominalCRE; Rebeiz *et al.* 2009). In Uganda, a partial selective sweep was detected at the latter element (Pool and Aquadro 2007; Rebeiz *et al.* 2009). This element drives expression of *ebony* in the abdomen that is repressed by an unknown enhancer in the intron (Rebeiz *et al.* 2009).

### *Soft selective sweeps can generate the pattern of polymorphism found at* ebony

The recurring pattern of isolated SNPs with high *F_ST_* raises the question of whether these data result from genetic drift and sampling variance, weak hard sweeps, or soft sweeps. We therefore performed a simulation analysis to identify evolutionary scenarios that might be consistent with the differentiation patterns revealed at melanin-synthesis genes in Ethiopia, *i.e.* low window *F_ST_* across these gene regions, and yet small numbers of SNPs showing strong population frequency differences. We focused on the case of *ebony* in Ethiopia, where one SNP was found to have *F_ST_* = 0.85, but a 5 kb window around this site had an overall Ethiopia-Zambia *F_ST_* of just 0.17.

In order to identify a plausible null model, we used δaδi (Gutenkunst *et al.* 2009) to estimate demographic parameters under a three population model, using allele frequency data from the middles of autosomal short introns (Halligan and Keightley 2006), from our Ethiopia, Rwanda, and Zambia populations. Parameter estimates, as detailed in Table S5, entailed a recent split between the Ethiopia and Rwanda populations (roughly 1,100 years ago, 95% CI 435 to 1,765), with an initial bottleneck to approximately 1% of the ancestral population size followed by exponential growth. Although we could not investigate all possible historical models, and the effects of natural selection on genomic variation could impact the precision of these estimates, the demographic estimates obtained here serve our primary objective of providing a neutral model capable of generating population genetic data resembling that of our empirical populations.

We then generated neutral simulation data to study window *F_ST_* and maximum SNP *F_ST_* under this null model. Compared with neutral simulations based on this demographic model, we found that our window *F_ST_* observed at *ebony* was only marginally elevated (*Q* = 0.07), but maximum SNP *F_ST_* deviated more strongly from neutral predictions (*Q* < 0.01; Figure 5). Thus, it is unlikely to observe this SNP frequency difference under the neutral model, and yet the window signal for *F_ST_* is probably insufficient to be detected in a typical genome scan.

**Figure 5.**
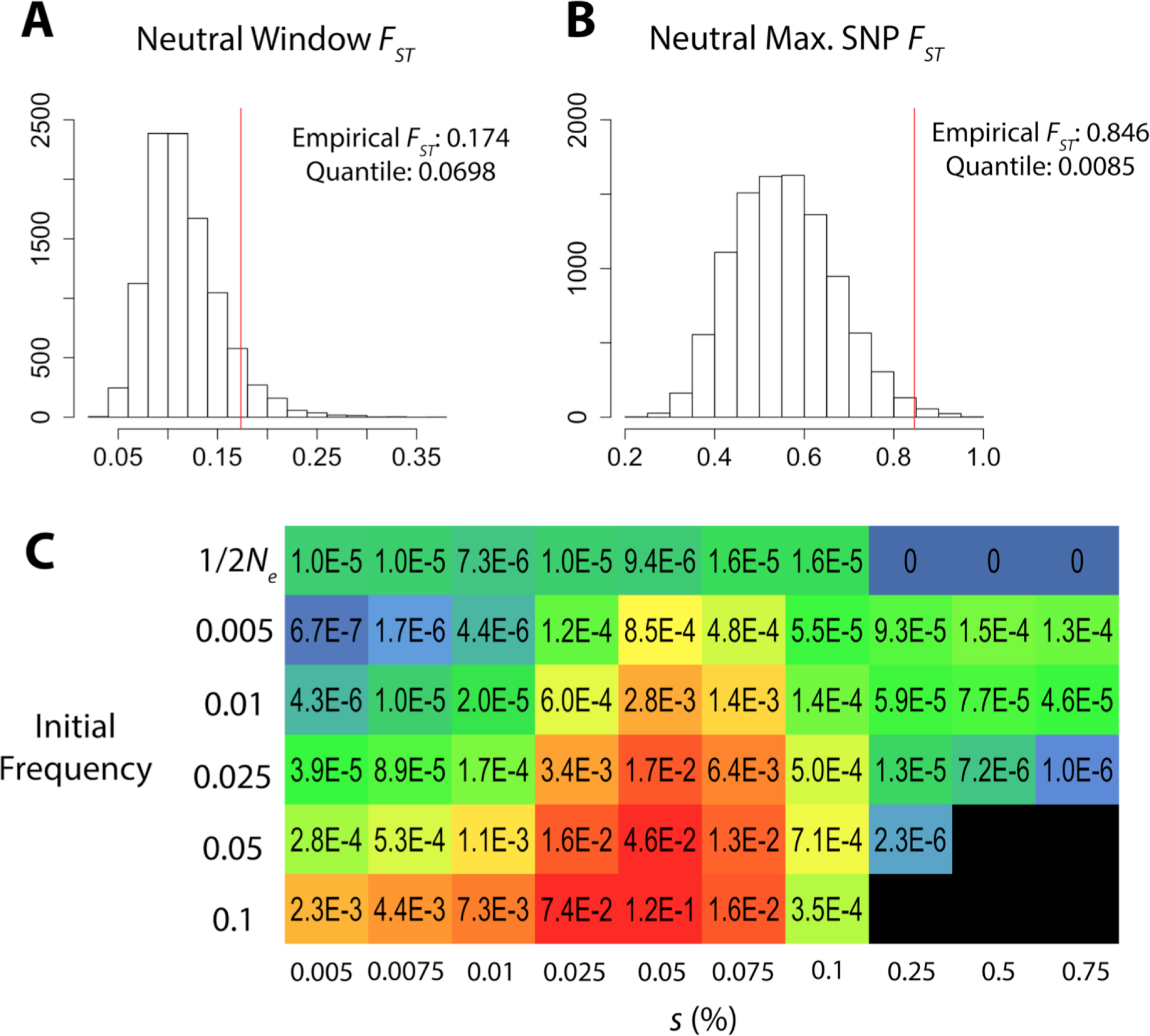
A simulation analysis was conducted to identify evolutionary models compatible with genetic differentiation at *ebony* between Ethiopia and Zambia populations. The top panels show that compared with neutral simulations, the empirically observed 5 kb window *F_st_* is only moderately elevated (A), but the maximum SNP *F_st_* observed at *ebony* is unusually high (B). The lower heat map (C) illustrates outcomes of simulations in which the most differentiated SNP was favored in Ethiopia. The acceptance rates depicted here depend on: (1) population allele frequencies at the focal SNP that are compatible with subsampling to match empirical counts, and (2) a window *Fst* at least as low as that observed at *ebony.* Acceptance rates are colored based on a log_10_ scale, with black cells indicating less than 10 successfully subsampled simulations out of 2.5 million. A range of selection strengths are depicted for models producing hard sweeps (initial frequency *1/2Ne)* and those conditioned on soft sweep outcomes (all others). Results suggest that soft sweep scenarios with higher initial frequencies are the most likely to raise the beneficial allele to high frequency (without fixing it), while also recapitulating the disparity between window *F_st_* and SNP *F_st_* observed at *ebony*.

Next, we conducted simulations based on the hypothesis that the most differentiated SNP at *ebony* was a target of positive selection in the Ethiopian population, varying the selection coefficient and the initial frequency of the beneficial allele. We rejected simulation replicates that could not be subsampled to match empirically observed allele counts for Ethiopia and Zambia at the target SNP (Materials and Methods). For each simulation scenario, we tested how often this SNP *F_ST_* was paired with a window *F_ST_* at least as low as we observed for *ebony.*

The scenarios that best matched the above criteria were soft sweeps with higher initial frequencies (Figure 5; Table S6). Weak hard sweeps could also match the *F_ST_* disparity, but these scenarios almost never allowed the beneficial allele to rise to high frequency on the time-scale of the Ethiopian population (Table S6). Hence, selection from standing genetic variation appears to be the strongest hypothesis for the patterns of genetic variation observed at *ebony* in the Ethiopian population.

## Discussion

We have integrated quantitative genetic and population genetic approaches to unravel the genetic basis of parallel melanism in three sub-Saharan populations of *Drosophila melanogaster.* Our BSA approach represents an unbiased genetic investigation of adaptive pigmentation variation, revealing between 1 and 10 QTLs per population/trait. The strongest QTL in each cross was estimated to explain 16% to 36% of the parental strain differences. Importantly, these effect size estimates can have upward bias in some scenarios, and should be viewed as preliminary (Pool 2016). Additional minor effect QTLs may exist below the threshold of our approach. Still, it seems clear that QTLs of moderate to large effect contribute to melanic evolution in these populations. Thus, in spite of its potentially polygenic architecture, our results indicate that the adaptive evolution of pigmentation involves a relatively modest number of loci, with some being unique and others being common among populations, supporting a convergent (yet dynamic) mode of oligogenic adaptation (Bell 2009; Martin and Orgogozo 2013).

Our QTLs showed an intriguing blend of parallelism and unpredictability within and between populations. In some cases, the same QTL peak was discovered from two independent crosses mapping the same trait in the same population, or across different populations (Table 1). In cases where the same locus may contribute to melanic evolution in two or more populations (i.e. Q1, Q7, Q10, Q11, Q14 and Q18), further study will be needed to assess whether the same variants may contribute in different populations, or if instead unique mutations at the same gene have given risen to similar phenotypes. As a potential example, Rebeiz *et al.* (2009) identified five causative cis-regulatory mutations at *ebony* underlying abdominal melanism in Uganda *D. melanogaster,* but although we also mapped a QTL to the *ebony* region in Ethiopia, previous analysis showed that a population from the Ethiopian highlands possessed only two of the mutations implicated in Uganda, and lacked the strongest contributor. Likewise, different mutations at the same genes drive parallel pigmentation changes in humans (Edwards *et al.* 2010; Yamaguchi *et al.* 2012) and closely-related species of *Peromyscus* mice (Manceau *et al.* 2011; Linnen *et al.* 2013). Our study does not confirm the precise molecular basis of pigmentation evolution in *D. melanogaster,* but it sets the stage for functional studies in these melanic populations.

When the same traits were mapped in separate crosses involving the same melanic population, some QTLs were identified repeatedly, but overall the results were quite variable (Table 1). At least for QTLs with effect sizes below 20%, one potential explanation is simply chance due to incomplete detection power. But in seven cases, stronger QTLs (bold in Table 1) were not replicated in another cross for the same population and phenotype, and these missing QTLs seem unlikely to result from type II error (Pool 2016). These results might stem from causative pigmentation variants that are not fixed differences between dark and light populations. Although the most differentiated SNPs at *ebony, tan, yellow,* and *black* are simply hypotheses for the mutational targets of selection, it is worth noting that none of these SNPs are fixed in Ethiopia, and only in the case of *tan* is the Ethiopian allele absent from our Zambia sample. Additional unpredictability in QTL mapping results could result if epistatic interactions contribute to pigmentation phenotypes. In other words, a given variant might have a strong detectable effect or a weak undetectable effect on a pigmentation trait, depending on the genetic backgrounds of the light and dark parental strains used in each mapping experiment.

We also showed evidence that melanic adaptation has likely involved selection on standing genetic variation. For Ethiopian abdominal background color, strong QTL peaks were observed near the pigmentation genes *ebony, tan, yellow,* and *black.* None of these loci showed a clear window signal of high genetic differentiation. And yet, it seemed unlikely that melanic evolution was instead caused by genes without known pigmentation functions that just happened to occur close to all of these pigmentation pathway genes. We therefore examined genetic differentiation at these genes more carefully, and found a recurring pattern of strong frequency differences limited to just one or a few SNPs. In the example of *ebony* in Ethiopia, simulations confirmed that this pattern seems best explained by selection on standing genetic variation. This conclusion mirrors the findings of other studies on the genetics of adaptation (Messer and Petrov 2013). For example, in spite of strong evidence for the anti-predatory adaptive significance of white coat in beach mice, no clear signature of selection was detected at the underlying *MCS1r* gene (Domingues *et al.* 2012). The potential importance of soft sweeps in adaptation to novel environments has significant implications for the population genetic identification of loci subject to local adaptation. While selection on relatively rare variants may lead to hard sweeps (Jensen 2014), at the other extreme, selection favoring alleles with appreciable prior frequencies may elevate multiple haplotypes, and the resulting genetic differentiation may be largely limited to the causative variant. In general then, it may be important to include single SNP analyses in scans for local adaptation, and not to rely solely upon window scans of genetic differentiation. Especially if applied to broad genomic regions, SNP-based approaches are likely to require larger sample sizes to avoid false positives due to random sampling variance.

Our work provides a case study of the genetic architecture of adaptive trait evolution, while providing a foundation for detailed molecular studies to confirm the relevant genes and mutations responsible for parallel melanic evolution in *D. melanogaster.* These results provide one partial set of answers to the basic questions posed at the beginning of this article concerning the genetics of adaptation. However, it will be important to conduct similar studies of different adaptive traits in different organisms to assess the generality of our findings.

Overall, our work highlights significant challenges to mapping the genetic basis of adaptive population differences. Even when relatively large effect loci are present, their detection may depend upon the specific genetic backgrounds used for QTL mapping. And genetic targets of local adaptation may not be detected in typical population genetic scans. Nevertheless, continued advances in the analysis of genomic data and molecular methods for confirming adaptive variants have the potential to expand our understanding of the nuanced process of adaptive trait evolution.

## Acknowledgments

This work was funded by NSF grant DEB-1049777 and NIH grant R01 GM111797 to JEP.

